# Differential sperm motility mediates the sex ratio drive shaping mouse sex chromosome evolution

**DOI:** 10.1101/649707

**Authors:** CC Rathje, EEP Johnson, D Drage, C Patinioti, G Silvestri, NA Affara, C Ialy-Radio, J Cocquet, BM Skinner, PJI Ellis

## Abstract

**Summary:** The search for morphological or physiological differences between X- and Y-bearing mammalian sperm has provoked controversy for decades. Many potential differences have been proposed, but none validated, while accumulating understanding of syncytial sperm development has cast doubt on whether such differences are possible even in principle. We present the first ever mammalian experimental model to trace a direct link from a measurable physiological difference between X- and Y-bearing sperm to the resulting skewed sex ratio. We show that in mice with deletions on chromosome Yq, birth sex ratio distortion is due to a relatively greater motility of X-bearing sperm, and not to any aspect of sperm/egg interaction. Moreover, the morphological distortion caused by Yq deletion is more severe in Y-bearing sperm, providing a potential hydrodynamic basis for the altered motility. This reinforces a growing body of work indicating that sperm haploid selection is an important and underappreciated evolutionary force.

## Introduction

The mouse Y chromosome, and to a lesser extent the X chromosome, exhibit an extraordinary level of gene amplification selectively affecting genes expressed in post-meiotic spermatids [1,2]. This is ascribed to a rampant and ongoing genomic conflict between the X and Y chromosomes over the control of offspring sex ratio. Partial deletions of the long arm of the Y chromosome (Yq) lead to sperm head malformation, sperm DNA damage, overexpression of sex-linked genes in spermatids and either a distorted offspring sex ratio in favour of females (for smaller deletions) or sterility (for larger deletions) [3–7]. Understanding the molecular basis of this sex ratio skew will answer a profound mystery -how genes break the first law of Mendelian genetics - and help develop models of sex chromosome differentiation, decay and conflict. Such knowledge may potentially also provide the basis for novel methods of sex selection in livestock animals.

As yet, the root cause of the skewing remains unknown. However, it is known to be regulated by a conflict between the sex-linked ampliconic transcriptional regulators *Slx* and *Sly* in haploid spermatids [8–12]. The prevailing model is that one or more X-linked genes acts post-meiotically to favour the production of female offspring, and that these genes in turn are repressed by (Y-linked) *Sly* and activated by (X-linked) *Slx*. Competition between *Slx* and *Sly* has led to a runaway “arms race” between the chromosomes that drives massive amplification of competing X / Y gene complexes, and also of any autosomal loci embroiled in the conflict. The signs of this conflict have now been observed at the molecular genetic level in the form of opposing transcriptional regulatory consequences of *Slx* / *Sly* depletion [4,9–11,13,14]; at the cellular biological level in the form of *Slx* / *Sly* competition for intracellular binding targets [14–17]; at the organism level in the form of sex ratio skewing in Yq-deleted animals, transgenic *Slx* / *Sly* knockdown animals and various Y chromosome congenic males [3,5,6,11,18]; at the ecological level in the form of population sex ratio skewing in introgression zones between subspecies with different copy numbers of the Y-borne amplicons [19,20]; and at the evolutionary population genetic level in the form of correlated co-amplification of gene families and recurrent selective sweeps at X and Y-linked amplicons [13,21–25].

Since *Sly* and *Slx* act as a “thermostat” affecting <u>*all*</u> transcriptionally-active sex-linked genes in spermatids, it is not currently possible to determine which of the hundreds of genes regulated by *Slx* / *Sly* are causative for the sex ratio skew, and which are merely “innocent bystanders” caught up in the feud [23]. Further insight into the paternal control of offspring sex ratio therefore requires identification of the physiological mechanism of the skew. Previous efforts in this direction have concentrated on identifying differences between Yq-deleted males and normal controls, revealing defects in acrosome biogenesis, sperm morphology and motility, *in vitro* fertilising ability, chromatin condensation of epididymal sperm, protease activity in the sperm head, and curiously an increase in progesterone production in the cumulus cells of daughters of affected males [3,5–7,26–35]. However, these analyses only give information about the <u>*global*</u> effects of Yq gene deficiency, and do not address the issue of <u>*differential*</u> effects on X- and Y-bearing sperm that mediate sex ratio skewing. Moreover, the majority of these studies have been performed in B10.BR-Y^del^ males that show very substantially reduced fertility in addition to sex ratio skewing [32]. It is therefore unclear which of the observed phenotypes relate to the sex ratio skew versus the fertility deficit, if indeed these are separable phenotypes.

The goal of the present study was thus to systematically identify the physiological differences between X- and Y-bearing sperm from Yq-deleted males that mediate the offspring sex ratio skew. We focused on the XY^RIII^qdel line that has a deletion of approximately ⅔ of Yq, since this has the largest known sex ratio skew of any Yq-deleted line with only minor accompanying fertility problems. Previous work has shown that across multiple different strain backgrounds, both inbred and outbred, the Y^RIII^qdel deletion leads to a six to ten percentage point deviation in offspring sex ratio in favour of females, relative to the strain background sex ratio [3]. In IVF experiments, we tested whether X- and Y-sperm from XY^RIII^qdel males have intrinsically different fertilising ability and/or differential ability to penetrate the cumulus surrounding the oocyte. Subsequently, we used newly-developed image analysis tools to search for morphological differences between X- and Y-bearing sperm, and *in vitro* swim-up testing to search for motility differences between X- and Y-bearing sperm. For the morphological analysis we also analysed shSLY knockdown males, which carry a short hairpin RNA construct that reduces *Sly* gene expression. shSLY males exhibit a similar degree of offspring sex ratio skewing to Yq-deleted males, but a much higher level of severe sperm malformation and impaired fertility [9]. This allowed us to explicitly test the relationship between specific categories of sperm malformation and sex ratio skewing. Importantly, shSLY males have no loss of Y chromosomal DNA, allowing us to test whether any morphological differences between X and Y-bearing sperm are a consequence of the reduction in DNA content in the Yq-deleted sperm.

## Results

### IVF abolishes offspring sex ratio skewing for MF1-XY^RIII^qdel males

In XY^RIII^qdel males, Ward and Burgoyne [36] showed that intracytoplasmic sperm injection (ICSI), which bypasses many aspects of fertilisation including cumulus and zona penetration, abolishes the offspring sex ratio skewing. Aranha and Martin-DeLeon [37] have also observed offspring sex ratio skewing in males heterozygous for Robertsonian (Rb) fusions involving chromosome 6, with a selective deficiency of female offspring carrying the fusion chromosome. In these males, the fusion chromosome bears inactivating mutations in the sperm head hyaluronidase *Spam1* [38–40] that are believed to be responsible for the skewed transmission. Thus, our initial hypothesis was that disruption of sperm/egg interactions - and in particular an interaction between sperm sex chromosome complement and hyaluronidase activity - might form the basis of the sex ratio skew.

For these embryo experiments, XY^RIII^qdel and control XY^RIII^ animals were maintained on an outbred MF1 genetic background. In our colonies, on this genetic background, the sex ratio for naturally-mated animals is 47.2% females for control XY^RIII^ males and 54.9% females for XY^RIII^qdel males, i.e. a 7.7 percentage point difference, similar to that seen in previous work [3]. First, we introduced an X-linked GFP transgene [41] onto the XY^RIII^qdel background to allow genotyping of preimplantation embryos. GFP transcription is visible from late morula stage onwards specifically in X^GFP^-positive embryos. Using sperm from the resulting X^GFP^Y^RIII^qdel males, we generated matched sets of IVF offspring with and without hyaluronidase pre-treatment of the oocytes to remove cumulus cells (n=6 males tested). Successfully fertilised embryos were cultured until blastocyst stage, and GFP expression scored in the inner cell mass by fluorescence microscopy. Contrary to our hypothesis that the sex ratio skew would depend on the presence of cumulus cells, the expected female skew was abolished in both the cumulus-on and cumulus-off experimental groups, with both instead showing a slight male bias consistent with the strain background sex ratio (Table 1, Figure 1).

**Table 1.**
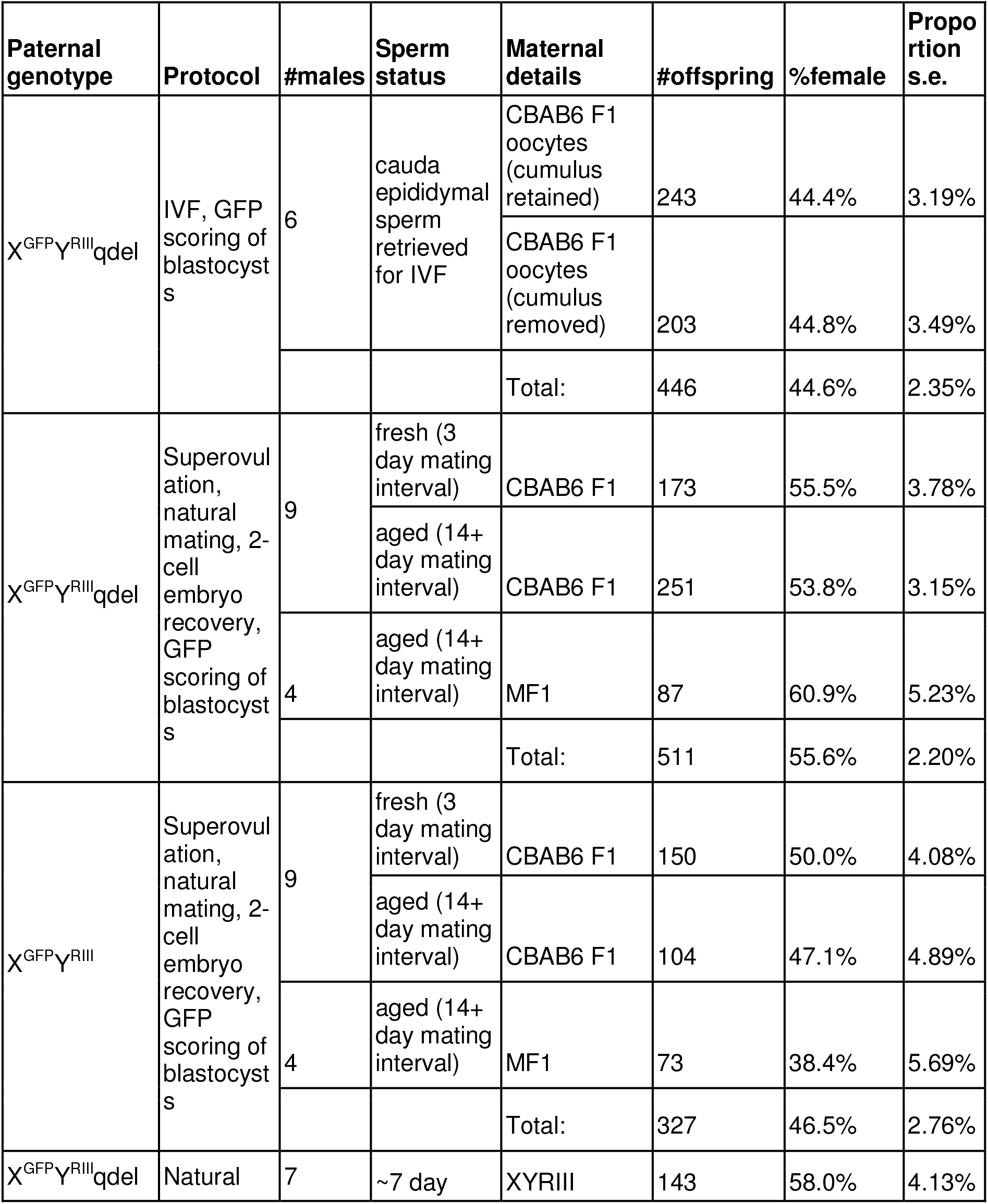
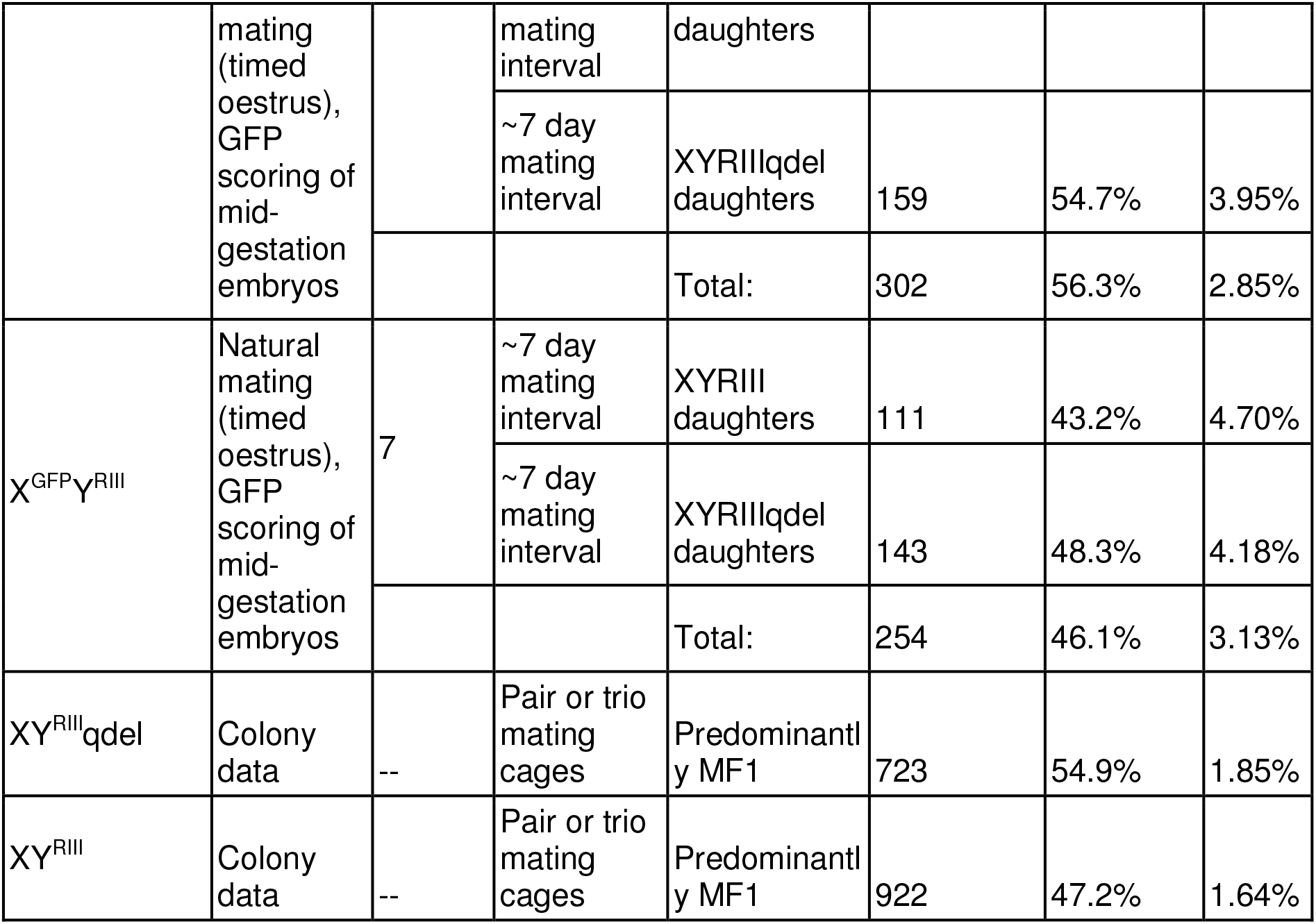
Offspring sex ratios shift in favour of females under natural mating in animals carrying the Y^RIII^qdel chromosome compared to animals carrying the Y^RIII^ chromosome when using both fresh and aged sperm; this difference is abolished by IVF using embryos with or without cumulus cells (see also Figure 1). S.E.P, standard error of proportion.

**Figure 1.**
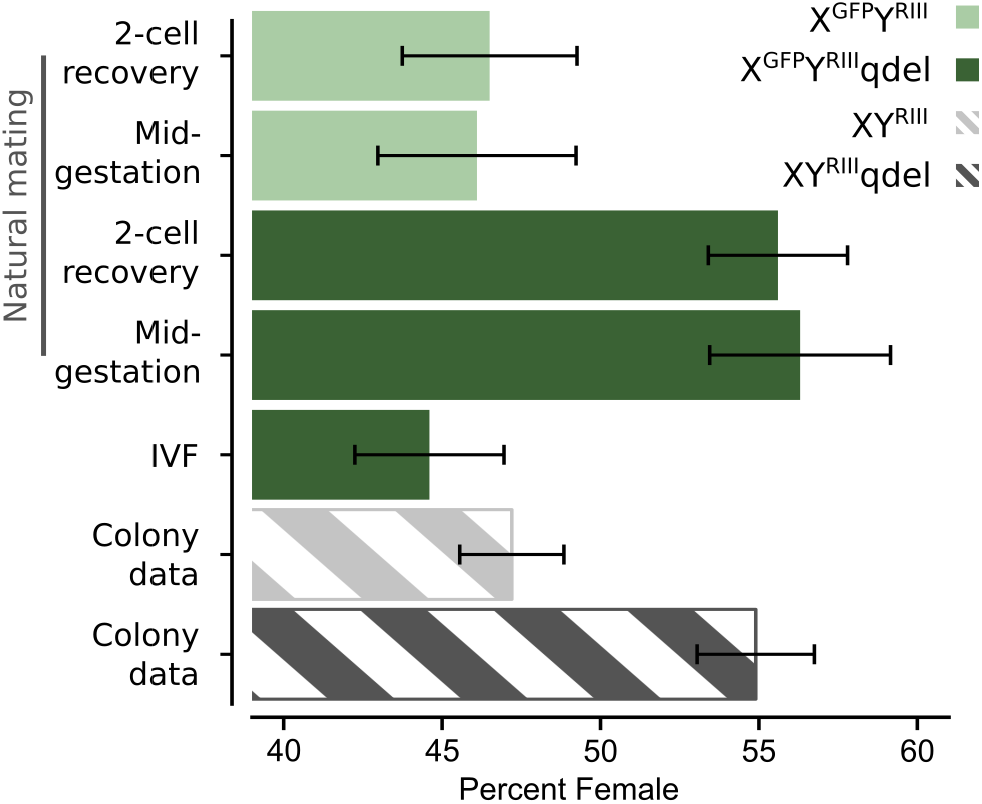
Sex ratios observed in colony mating and in embryos generated by IVF versus natural mating and scored at differing time points, see also Table 1. XY^RIII^qdel animals and X^GFP^Y^RIII^qdel embryos show a marked sex ratio skew in favour of females when mated naturally, but this is abolished in IVF experiments. XY^RIII^qdel animals, X^GFP^Y^RIII^ and IVF-derived X^GFP^Y^RIII^qdel embryos show a slight skew in favour of males. Error bars show standard error of proportion.

Next, to test whether the abolition of the sex ratio skew in the IVF cohorts was due to the fertilisation procedure, and not an artifact of the embryo culture step, we mated X^GFP^Y^RIII^qdel and control X^GFP^Y^RIII^ males (n=9 males tested per genotype) to superovulated females to allow natural fertilisation, collected the resulting embryos at the 2-cell stage by flushing oviducts at 1.5 days *post coitus*, and cultured them *in vitro* to blastocyst stage for GFP scoring. Since the transmission skew in chr. 6 Robertsonian fusion heterozygous carriers is reduced when sperm are subjected to prolonged aging in the epididymis [42], in this natural mating / embryo recovery experiment we also took the opportunity to test whether the sex ratio skew was affected by epididymal aging in either X^GFP^Y^RIII^qdel or control males, by using inter-mating intervals of 3 days for fresh sperm versus 14 days for aged sperm. We observed no difference between fresh and aged sperm for either genotype, but there was a significant skew in favour of females in the offspring of X^GFP^Y^RIII^qdel males (combining aged and fresh sperm data, two-tailed binomial p=0.0113 relative to null expectation of 50:50 ratio, p=0.0102 relative to MF1-X^GFP^Y^RIII^ control data, p=0.0007 relative to MF1-X^GFP^Y^RIII^qdel IVF data, Table 1).

Finally, given the documented imprinted effects of Yq deletion on the cumulus cell properties of daughters of B10.BR-Y^del^ males [27], we tested whether there was an imprinted effect on the sex ratio skew by mating X^GFP^Y^RIII^qdel and X^GFP^Y^RIII^ males (n=7 males tested per genotype) to daughters of XY^RIII^qdel or XY^RIII^ males and scoring the resulting offspring. In this experiment, the females were not superovulated, and the embryos were dissected and scored for GFP in mid-gestation. While there was again a significant skew towards females in the offspring of X^GFP^Y^RIII^qdel males (two-tailed binomial p=0.0285 relative to null expectation of 50:50 ratio, p=0.0162 relative to MF1-X^GFP^Y^RIII^ control males), there was no effect of the maternal background. As a control for unexpected effects of the X^GFP^ transgene, we compared the results from our X^GFP^Y^RIII^qdel and X^GFP^Y^RIII^ embryo studies to colony breeding data for the co-located XY^RIII^qdel and XY^RIII^ colonies (i.e. the identical genetic backgrounds with no GFP transgene), which showed that there was no effect of the X-linked GFP on offspring sex ratio. The slight male bias seen in the X^GFP^Y^RIII^qdel IVF embryos was indistinguishable from the slight male bias characteristic of the overall strain background.

Collectively, these *in vivo* and *in vitro* fertilisation experiments demonstrate that the sex ratio skew in the offspring of Yq-deleted males is evoked specifically during natural mating/fertilisation, is abolished by IVF, is not modified by the epididymal transit time of the sperm, and is not affected by the maternal genetic backgrounds tested. We conclude that the mechanism of the skew cannot be related to any of cumulus penetration, zona pellucida or oolemma binding, sperm/egg fusion or subsequent embryonic development, but must relate to transport of the sperm to the site of fertilisation.

### Yq deletion affects Y-bearing sperm morphology more severely than X-bearing sperm

Given the above, since the female tract has been shown to discriminate between morphologically normal B10.BR sperm versus abnormal B10.BR-Y^del^ sperm at the uterotubular junction [43], we considered that this might lead to sex ratio skewing if Y-bearing sperm are more severely morphologically distorted than X-bearing sperm in Yq-deleted males. We therefore systematically tested for morphological differences between X- and Y-bearing sperm, using a novel image analysis tool for quantitative sperm morphometry. The design and validation of this software has been described in recent publications [44,45]. In order to compare X- and Y-bearing sperm, we used a repeat-imaging protocol to measure cells both before and after FISH hybridisation. This allowed us to capture pre-FISH morphology data and correlate this with post-FISH identification of X-versus Y-bearing status (**Figure S1**).

In this experiment we analysed the effect of Yq deletion on both MF1 and C57Bl6 genetic backgrounds. This allowed us to also analyse shSLY males, which are maintained on a C57Bl6 background. Colony breeding data showed that on a C57Bl6 genetic background, control XY^RIII^ males produce 51.66% female offspring, while XY^RIII^qdel males produce 61.36% female offspring. This is a 9.7 percentage point difference, i.e. although the MF1 and C57Bl6 colonies have a different underlying wild type sex ratio, Yq deletion leads to a similar effect size on both genetic backgrounds. Only two litters’ data were available for shSLY males (14 female and 2 male pups). This is significantly different from a 50:50 ratio, confirming that shSLY knockdown leads to sex ratio skewing on a C57Bl6 background, but the sample size is insufficient to permit a reliable estimate of the magnitude of the skew. On both genetic backgrounds, our morphological analysis confirmed the known manifestations of Yq deletion including increased acrosomal curvature, and shortening of the apical hook (Figure 2A,B). Surprisingly, both Yq-deleted sperm and shSLY sperm also showed a marked reduction in all linear dimension measures and in overall cross-sectional area relative to wild type sperm, which has not been previously described (Figure 2C).

**Figure 2.**
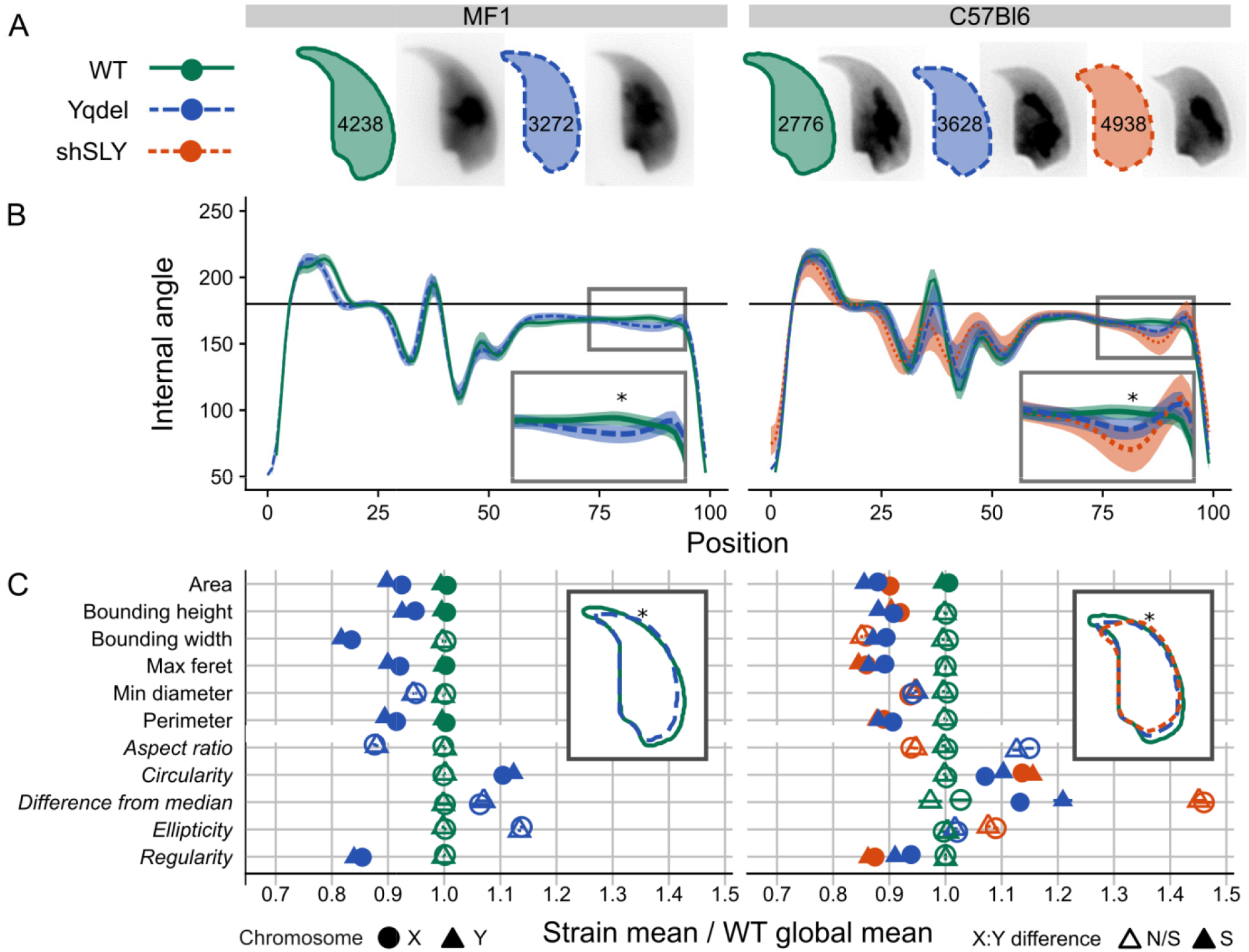
A) Consensus nuclear outlines for each strain alongside an example DAPI-stained nucleus. Numbers indicate the number of nuclei analysed for each strain. B) Angle profiles from each strain [44]. The X axis is an index representing percentage of the total perimeter as measured counterclockwise from the apex of the sperm hook. The Y axis represents the interior angle measured across a sliding window centered on each index location - a smaller angle represents sharper curvature at any given point, thus (e.g.) the hook apex at index 0 shows the smallest angle. The insets highlight the increased acrosomal curvature seen in the mutant genotypes, with ‘*’ indicating the point of greatest difference in curvature between genotypes, at index 85. C) Comparison of X- and Y-bearing sperm in standard morphometric parameters (see [44] for definitions), compared to mean wild-type values. Italics indicate size-independent parameters. Inset: overlapping consensus nuclear outlines for each genotype, with the location of index 85 marked ‘*’.

For the majority of parameters tested, including both size-dependent measurements (linear dimensions and area) and size-independent measurements (circularity and regularity), Y-bearing sperm in both mutant genotypes were significantly different from X-bearing sperm, and deviated more from wild-type values. We therefore tested explicitly whether there was any difference in phenotype severity between X- and Y-bearing sperm in each genotype for the three main manifestations of the Yq deletion phenotype: acrosomal curvature, hook shortening and area reduction. In Yqdel males, but not in wild type or shSLY males, the degree of acrosomal curvature (Figure 3A) was significantly more severe in Y-bearing sperm than in X-bearing sperm (Mann-Whitney tests, p<0.001). Similarly, in Yqdel males, hook length was shorter in Y-bearing sperm than X-bearing sperm (data not shown). We were unable to analyse hook length in shSLY sperm since a high proportion of sperm were so strongly distorted that hook length could not be clearly defined.

**Figure 3.**
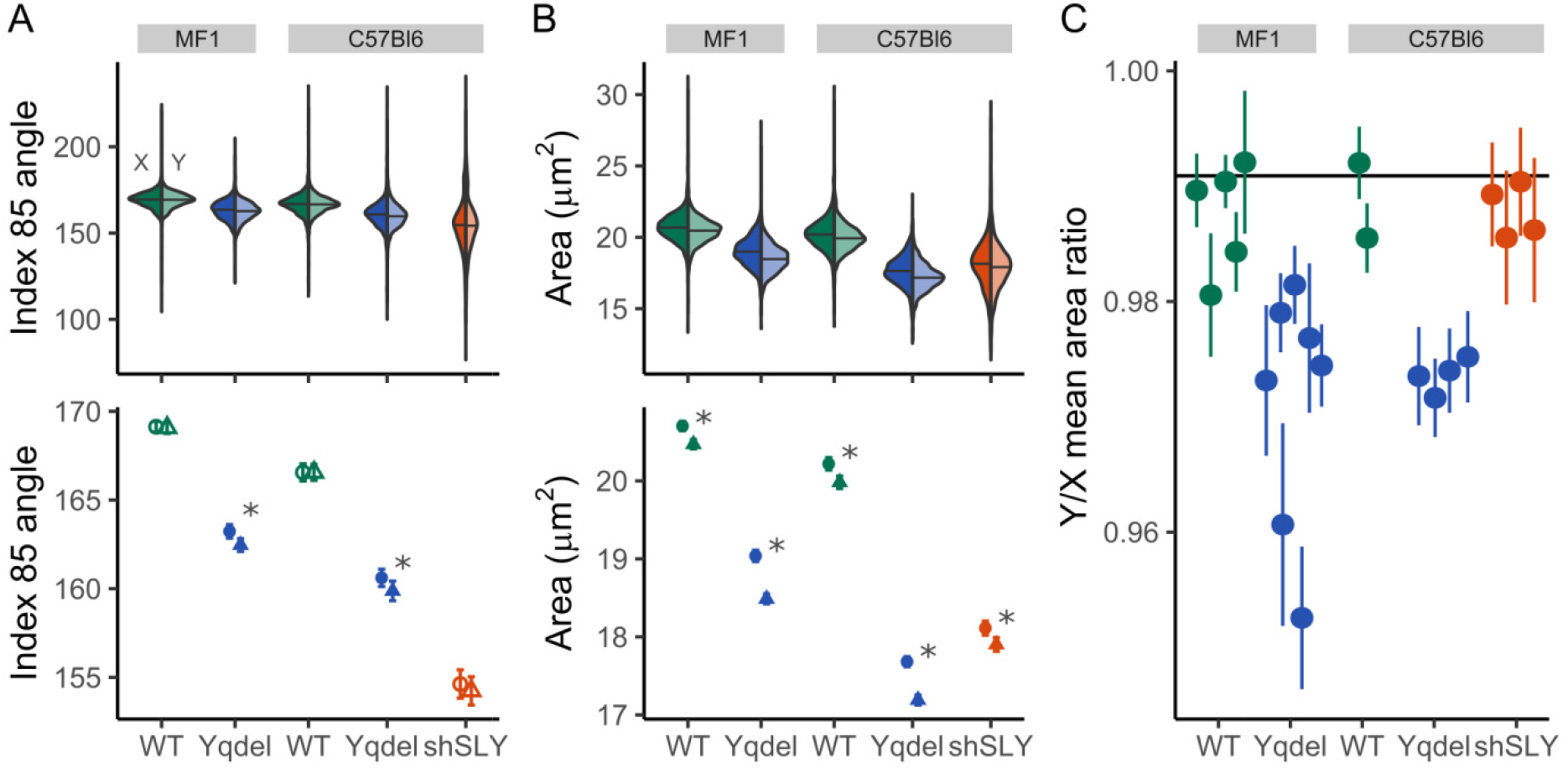
A) Angles at profile index 85 in each strain, comparing X- and Y-bearing sperm. The upper panel shows violin plots of the complete population data for each strain, while the lower panel shows the mean and standard error of the mean in order to highlight differences. Significant XY differences (Mann-Whitney test, p < 0.05) are marked ‘*’. Y-bearing sperm have a significantly more curved acrosome in Yqdel animals only. B) Comparison of nuclear areas, using the same format as (A); Y-bearing sperm are consistently smaller than X-bearing sperm in all genotypes including wild types. C) Comparison of the mean Y/X area ratio in each sample (see **Table S1** for the details of sample numbers analysed per genotype), with standard error of the mean. The horizontal line indicates the mean wild type ratio. Wild type and shSLY samples have similar area ratios, while Yqdel samples show a further reduction in Y area.

Since sperm head area showed a significant difference in wild type as well as mutant sperm on both genetic backgrounds, we considered this was likely to be affected by the haploid genome size. Quantifying the differences in each genotype, we found that irrespective of genetic background, Y-bearing cells from the WT and shSLY genotypes were on average 1.1%-1.3% smaller in cross-sectional area than X-bearing sperm (Figure 3B,C; **Table S1**). Assuming an equivalent reduction in thickness, this corresponds to an estimated 1.7%-1.8% reduction in nuclear volume, around half that expected given the known 3.2% difference in DNA content as measured by flow cytometry [46] and calculated from the known genomic sizes of the X and Y chromosomes (i.e. an 80 Mb difference compared to a 2.6Gb haploid genome). In Yqdel males, on both genetic backgrounds the Y-bearing sperm were on average 2.7%-2.8% smaller in cross-sectional area than X-bearing sperm. This corresponds to an estimated 4.1% difference in nuclear volume between Y-bearing sperm and X-bearing sperm. Thus, an additional 2.3%-2.4% difference between X and Y-bearing sperm volume is ascribable to the effects of Y chromosome deletion. This corresponds almost perfectly to the estimated 2.3% difference in DNA content between sperm bearing a normal versus a Yq-deleted Y chromosome (i.e. a 60 Mb deletion compared to a 2.6Gb haploid genome). We conclude that in animals with a normal karyotype, sperm size is “buffered” such that sperm are more similar in volume than their DNA content would indicate. However, pathological genomic deletions are not “buffered” and so sperm size scales linearly with the amount of DNA loss.

### Cluster analysis reveals an X/Y gradient of morphological abnormalities

Following this morphometric analysis of sperm dimensions, we carried out dimensional reduction on size-independent angle profiles using t-SNE, and used hierarchical clustering to identify different sperm head shape groups within each genetic background. On the MF1 strain background this separated sperm shapes into two major clusters, the first containing predominantly but not exclusively XY^RIII^ sperm and the second containing predominantly but not exclusively XY^RIII^Yqdel sperm. This could then be subdivided into grades of abnormality, i.e. clusters N1/N2 and A1/A2. As expected from the dimensional measurements, the more severely affected sperm heads from XY^RIII^Yqdel males were preferentially Y-bearing cells, while the more morphologically normal sperm heads were preferentially X-bearing cells (Figure 4A,C; Table 2).

**Table 2.**
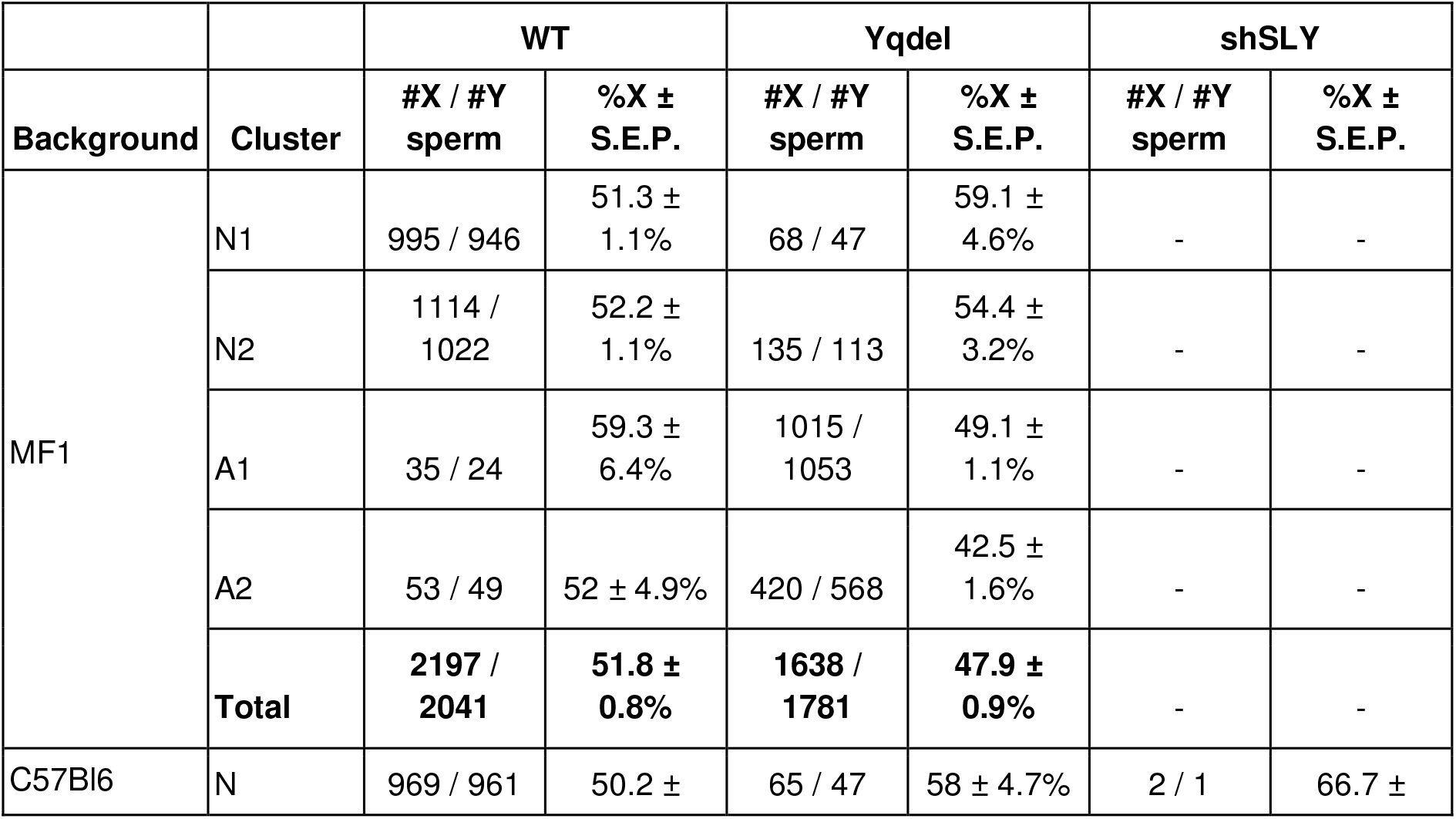
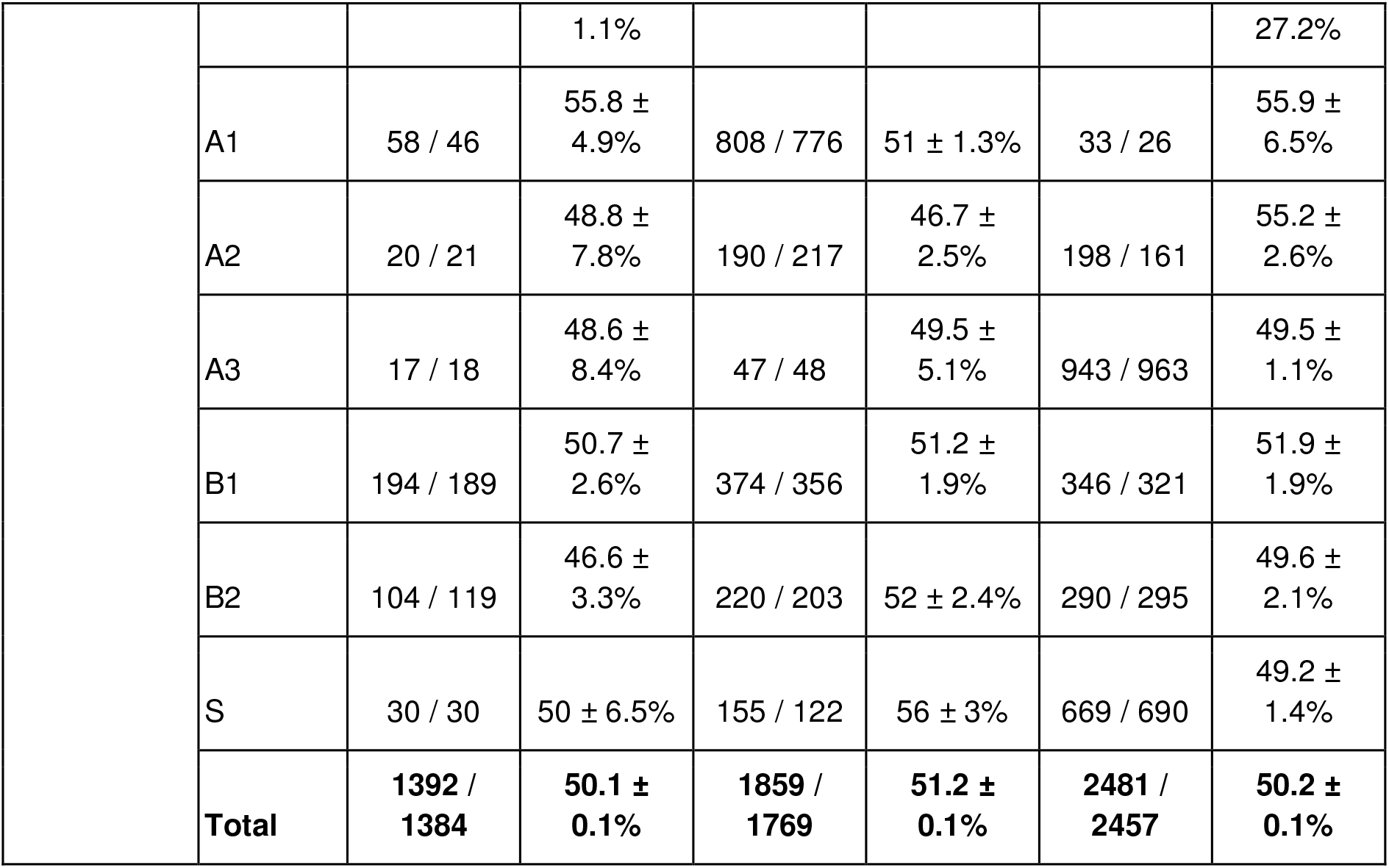
Number and proportion of X- and Y-bearing sperm in each morphological cluster (see also Figure 4). On an MF1 background, more abnormal sperm are enriched for Y-bearing sperm (Fisher’s exact test, p=0.002) in Yqdel samples, but not in WT samples (p=1). On the C57Bl6 background, a similar gradient is observed in Yqdel and shSLY samples, but is not statistically significant in these (p=0.63, 0.66), or in WT (p=1). S.E.P, standard error of proportion.

**Figure 4.**
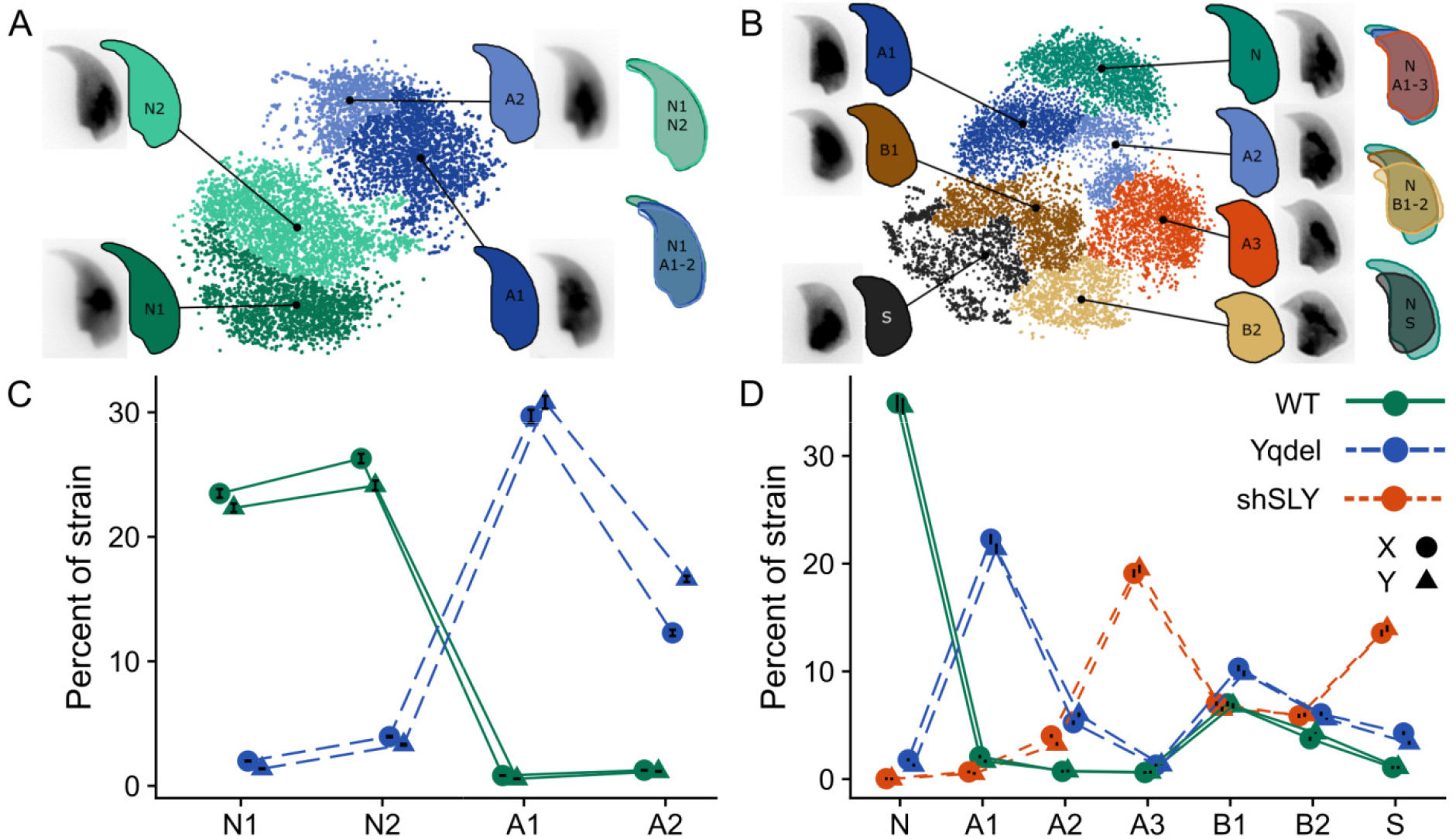
A,B) Clustered t-SNE plots of angle profiles from MF1 and C57 samples respectively allow groups of nuclei with similar shapes to be distinguished. Consensus nuclear outlines are shown for each cluster alongside a representative DAPI-stained nucleus. Overlapping consensus nuclei allow comparison of clusters to the most ‘normal’ category. C,D) The proportion of sperm in each strain within each cluster are shown, separated into X- and Y-bearing groups. Error bars show the standard error of proportion. More clusters with severe ab ormalities are enriched for Y-bearing sperm in both backgrounds (see also Table 2); C57Bl6 genetic background has additional shape abnormalities (B1, B2) that are also present in wild type males and thus are not related to the Yq deficiency phenotype.

On the C57Bl6 strain background the picture was more complex. Once again there was a continuous spectrum of increasing shape abnormality (clusters A1/A2/A3) corresponding to increasing severity of the Yq gene deficiency phenotype of acrosomal curvature, hook shortening and area reduction. However this dataset additionally presented a second axis of shape abnormalities (clusters B1/B2) that encompassed shortening of the sperm nucleus, “flaring” of the base of the sperm head and exaggeration of the dorsal angle in addition to acrosomal curvature, hook shortening and area reduction. Since this second type of abnormality was also present at comparable levels in the WT controls, we interpret these as being a strain characteristic unrelated to the Yq gene deficiency phenotype. Finally, a third type of abnormality (cluster S), that encompassed narrowing of the base of the sperm head, almost total loss of acrosomal hook and effacement of the tail attachment site was almost exclusive to shSLY sperm.

Of these three types of abnormality, the first type (i.e. related to Yq gene deficiency) again showed an X/Y gradient in the affected cells, although this was not statistically significant (Figure 4B,D; Table 2). In particular, the number of sperm and the X/Y ratio in the two least morphologically abnormal categories (N and A1) was consistent with both the direction of sex ratio skew in XY^RIII^Yqdel and shSLY, and also the marked fertility reduction in shSLY males. Therefore, despite the confounding effects of unrelated background sperm morphology distortions, the cluster analysis overall supports the circularity and regularity measurements showing that on both genetic backgrounds, Yq deletion and shSLY knockdown lead to more severe morphological defects in Y-bearing sperm. On the C57Bl6 background, we also analysed chromosome localisation within the nucleus using dynamic warping [45] (**Figure S2**). This showed no change in chromosome territory localisation, indicating that the morphological changes are likely driven by changes in cytoskeletal dynamics during spermiogenesis rather than differences in chromatin organisation.

### Y-bearing sperm have lower motility than X-bearing sperm in XY^RIII^qdel males

Finally, we directly tested whether there are motility differences between X- and Y-bearing sperm in XY^RIII^qdel males, once again working on an MF1 strain background. While conventional computer aided semen analysis (CASA) has shown impairment in a range of motility parameters in B10-BR.Y^del^ sperm relative to B10.BR sperm [26,34], this method cannot distinguish between X- and Y-bearing sperm. We therefore devised a swim-up protocol to separate sperm into six fractions according to their relative motility (see Methods). As a control, the s me experiment was performed with sperm killed by repeated freeze-thaw cycling. As expected, the sperm count in the upper, more highly motile fractions was very greatly reduced in t e killed sperm samples, with only trace levels of contaminating cells remaining. We analysed 9 WT and 10 Yqdel males, and counted at least 400 cells from each male for each fraction for both live and killed cells to obtain an accurate measurement of the proportion of X- and Y-bearing sperm: a total of 91,015 cells for the dataset presented here (Figure 5).

**Figure 5.**
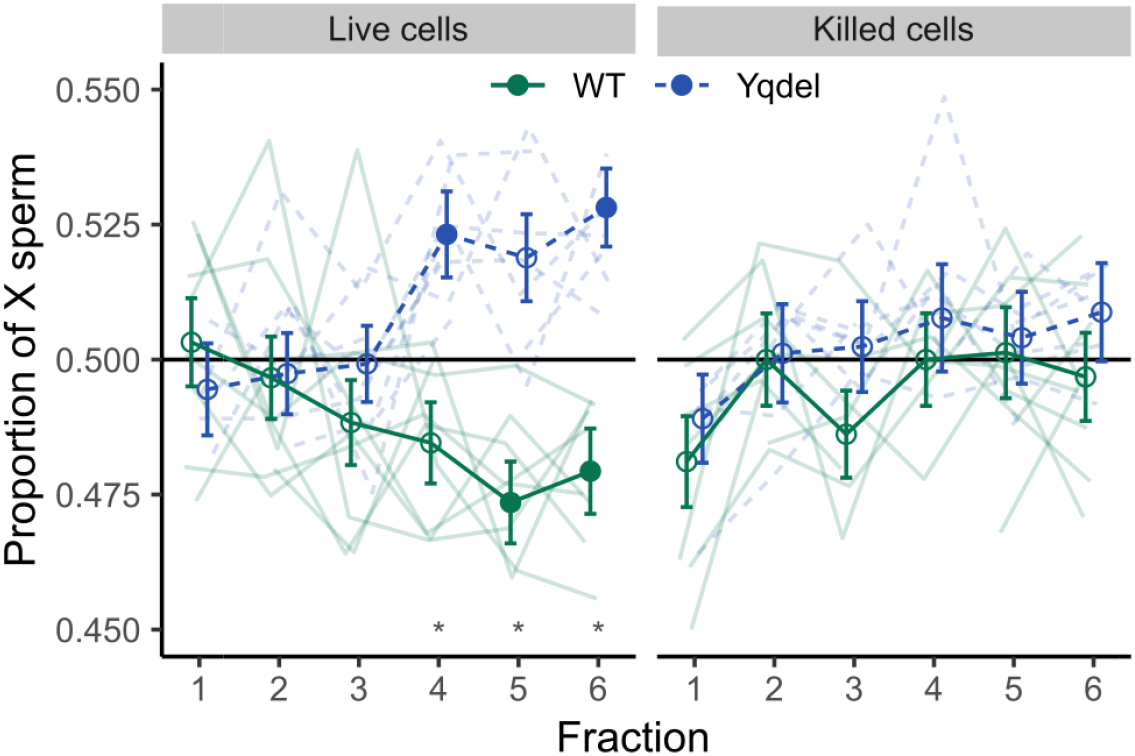
X-axis indicates sperm fractions with relatively increasing motility. Highly motile sperm are enriched for X-bearing over Y-bearing sperm in Yqdel samples, but the reverse in wild type samples. The negative control (freeze/thaw killed sperm) showed no enrichment for X or Y sperm in any fraction. Individual samples are shown in faded lines. Stronger lines show the means and standard errors of proportion per fraction per genotype. Closed symbols indicate fractions that were individually different from a 50:50 ratio, open symbols indicate fractions that are not significantly different from 50:50 (one sample Z-test, p<0.05 after Bonferroni-Holm multiple testing correction). ‘*’ indicates fractions where the difference between Yqdel and WT genotypes is significant (two sample Z-test, p<0.05 after Bonferroni-Holm multiple testing correction).

Observers were blinded to the live / killed status of sperm samples and also to the fraction analysed, i.e. blind to high / low motility status. Blinding by genotype was not possible since the shape difference between XY^RIII^qdel and XY^RIII^ sperm remains identifiable after FISH hybridisation. In both XY^RIII^qdel and XY^RIII^ samples, the lowest-motility live cells and all of the killed cells showed no significant deviation from a 50:50 ratio of X- and Y-bearing sperm. In the fractions with increasing motility, there was a progressively-widening and strongly significant difference in X:Y sperm ratio between the XY^RIII^qdel samples and the XY^RIII^ control samples, with the XY^RIII^qdel samples being enriched for X-bearing sperm and XY^RIII^ samples for Y-bearing sperm, consistent in direction with the observed sex ratios in colony breeding data for each line. Both enrichments were significantly different from a 50:50 ratio and from each other. Fitting a fully-specified beta regression model with a logit link (**Table S2**) yielded a likelihood ratio test p value of 2.43 × 10^−07^ against the null distribution, and a pseudo-R^2^ of 0.87, indicating that the model describes the data well. A p value of 3.26 × 10^−08^ for the triple interaction term (Fraction | Genotype | Live) indicates high confidence that there is a linear relationship between sperm sex ratio and motility, and that this is dependent upon both the genotype and live/dead status of the sperm samples.

In this experiment, the sperm sex ratio deviation, while consistent in sign, was lesser in magnitude than the sex ratio skew seen in the colony breeding data. This may be because performing the fractionation in a large volume measures increased diffusion via increased general motility, rather than progressive linear velocity. Another contributing factor is likely to be the absence of the various strictures in the female reproductive tract that select for highly motile versus weakly motile cells [47]. We note however that since the bottom, least motile fraction contains an equal proportion of X- and Y-bearing sperm, any contamination of the upper, more motile fractions necessarily serves to reduce the skew, and that our results are therefore a conservative measure of X / Y motility differences. We further looked for differences in sperm midpieces using MitoTracker staining. No MitoTracker staining was present in the killed sperm samples; in the live samples, the swim-up fractionation enriched for sperm with longer midpieces in both XY^RIII^ and XY^RIII^qdel genotypes (**Figure S3**). The XY^RIII^qdel sperm in general had poorly active mitochondria, but swim-up fractionation enriched for XY^RIII^qdel sperm with more active mitochondria. This therefore confirms that the fractionation procedure enriches for the sperm with the best motility-related mitochondrial parameters.

## Discussion

The work presented here is the first ever study to show a physiological difference between X- and Y-bearing sperm in a mammalian model and link this to the subsequent effects on offspring sex ratio. In the search for the specific genes causative for the sex ratio skew in XY^RIII^qdel males, our results open a new avenue of investigation by outlining the underlying basis of the phenotype: differential sperm motility, associated with differential morphological distortion of X- and Y-bearing cells. We cannot as yet say whether the changes in sperm shape lead directly to the altered motility by affecting hydrodynamic efficiency, or whether the sperm also have altered biomechanics (e.g. differential flagellar beat patterns as in the case of *t* heterozygotes [48] and/or biochemistry (e.g. differential capacitation or hyperactivation kinetics). Studying these aspects of the phenotype will require improved methods to separate sperm with differing motility and probe their biochemistry and chromosomal content, with recent advances in microfluidics providing a potential route forward [47]. This work adds to the growing body of experimental and theoretical work demonstrating that even in animal sperm that undergo postmeiotic transcriptional shutdown, haploid selection within a single ejaculate is an important and underappreciated evolutionary force [49–54]. Our findings are supported by over two decades’ worth of research by multiple groups that have demonstrated the evolutionary consequences of this sex ratio conflict for mouse genome structure and population genetics.

The question of whether there are morphological differences between X- and Y-bearing sperm has provoked controversy for decades [55–59]. Here, we identify a slight difference in size between wild type X and Y-bearing sperm nuclei, and show that a genomic deletion further reduces Y-bearing sperm size in Yqdel males. The baseline X/Y size difference is consistent between both MF1 and C57Bl6 genetic backgrounds, and between both wild type and shSLY genotypes, despite the different underlying sperm size in each. However, the magnitude of the X/Y difference is significantly lower than predicted from DNA content, implying that nuclear size is somehow adjusted (potentially by alterations in chromatin condensation) to reduce the size difference between X- and Y-bearing sperm from karyotypically normal animals. This is consistent with recent findings in cattle, which showed a 1.4% difference in sperm head area between X- and Y-bearing sperm. This corresponds to a 2.1% difference in sperm nuclear volume, as compared to a 4% difference in DNA content in cattle [60]. This “buffering” of X/Y sperm nucleus size differences may explain previous contradictory findings in this regard [57]. Moreover, the X/Y size difference in karyotypically normal animals is several times smaller than the within-strain coefficient of variation in both mouse (4.5-6.0%, our data) and cattle (6.66% [60]) and is only detectable by measuring large numbers of cells. Thus, it is futile to seek to sort X and Y-bearing sperm by physical size in individuals with a normal karyotype. Size-based selection might however in principle be able to enrich for normal versus abnormal sperm if large chromosomal aberrations are involved.

In addition to quantifying X/Y sperm size differences, we show that the morphological abnormalities induced by Yq deletion and/or shSLY knockdown are more severe in Y-bearing sperm than in X-bearing sperm on both genetic backgrounds, and that Y-bearing sperm from Yqdel males have relatively worse motility than X-bearing sperm on an MF1 background. Since the sex ratio skew is abolished by IVF, the morphological differences alone are not sufficient to alter sex ratio - i.e. all or most shapes of sperm must be competent to fertilise the oocyte, consistent with prior findings via ICSI in Balb/c wild type males [61] and in *Sly* knockdown males [12]. Rather, the morphological distortion must impair the sperm’s ability to transit from the site of deposition to the fertilisation site, potentially via altering the cell’s hydrodynamic properties. Since the degree of X sperm enrichment we obtained in our *in vitro* swim-up experiment was significantly less pronounced than the sex ratio difference seen *in vivo* during natural mating, it is likely that selection of the most highly motile sperm is more efficient in the environment of the female reproductive tract. We cannot however rule out the possibility that the female tract may also directly select X-versus Y-bearing sperm via (e.g.) surface antigenic differences. The difference in X-versus Y-bearing sperm velocity is likely to be small in absolute terms since the winner-takes-all nature of the race to fertilise the egg implies that small differences in average sperm velocity can lead to much larger changes in the proportion of X- and Y-bearing sperm in the vanguard population arriving at the site of fertilisation in the oviduct [50]. Put simply, Michael Phelps is the most-decorated Olympic swimmer of all time, despite only needing to be 1/1000 of a second faster than his nearest competitor over the course of a several-minute race.

The next challenge is to elucidate the causal chain between *Slx*/*Sly* genomic competition and the physiological differences identified in this study. In this endeavour, other examples of male-mediated transmission ratio distortion such as the Robertsonian chromosome fusion example discussed above, and the *t* complex autosomal drive system give a clue to potential molecular mechanisms. In each, there is at least one “responder” gene whose products escape sharing across the cytoplasmic bridges between syncytially developing spermatids [62–64]: *Spam1* in the case of the Rb fusions [38–40], *Smok^TCR^* in the case of the *t* complex [65,66]. An alternative is that there is no “responder” gene, and that some gene on the X chromosome is directly toxic to the Y-bearing cells, e.g. by targeting some Y-specific DNA element, as in the case of HP1D2 in *Drosophila* [67].

We observed sex ratio and sperm motility deviations in favour of males in the control MF1-XY^RIII^ population in this study. While this finding was unexpected, we note that this line carries a *Mus musculus musculus* Y chromosome (with a high copy number of Y-linked amplicons) on a mainly *Mus musculus domesticus* background (with a low copy number of X-linked amplicons) [11]. This is predicted to unbalance the genomic conflict in favour of producing more male offspring, as observed in previous work with a range of Y chromosome consomic lines on a C57Bl6 strain background; see figures 4B and 5A in [18], and in transgenic knockdown of *Slx* [10]. This kind of “thermostat”, which generates both male and female skews depending on the balance of X- and Y-linked amplicons, is hard to reconcile with a simple X-borne, DNA-directed toxin system. We therefore predict that there is likely to be a “responder” gene on the X or Y chromosomes that escapes sharing between syncytial spermatids and acts to “label” X- or Y-bearing sperm. Identification of this responder gene remains a significant goal of future work in this model.

A final important question is whether the genomic conflict underlying the sex ratio skew is mouse-specific, or whether it is present in other taxa including livestock species where the potential for sex ratio manipulation has considerable ethical and economic implications. While the upstream transcriptional regulators *Slx* and *Sly* are specific to the Palaearctic radiation of mouse species, their mode of action appears to involve interaction with *Ssty*, which is present more widely in rodents and has become independently amplified in rat, together with its X-linked homologue *Sstx* [13,68,69]. Together with data indicating the presence of paternally-mediated sex ratio skewing in an even more distantly related rodent species, *Peromyscus* [70], and the presence of sequences related to *Ssty* in the genomes of *Ellobius* species [71], this suggests that the genomic conflict may be much longer-standing. The repeated expansions of X and Y ampliconic genes in rodents may represent successive outbursts of conflict as new “control knobs” arise to regulate the underlying biology. Under this hypothesis, mouse *Slx* / *Sly* are simply the most recently-evolved participants in an overall layer cake of genomic warfare spanning at least 25 million years of evolution. This in turn may explain why *Slx* and *Sly* act via modulation of post-meiotic sex chromatin (PMSC), since this allows them to simultaneously regulate multiple linked drive genes on the sex chromosomes. A key question for future work is to trace the history of X and Y amplicon expansion, both within muroid rodents, but also in non-rodent mammalian species: This will allow identification of when the process started, which species it affects, and how and when different genes have arisen and become recruited into the conflict.

## Acknowledgments

We thank the animal handling staff at the University of Kent, University of Cambridge, Cochin Institute and Charles River Laboratories. B.M.S. was supported by the Biotechnology and Biological Sciences Research Council (BBSRC, BB/N000129/1). P.E. and C.C.R. were supported by H.E.F.C.E. (University of Kent) and by the BBSRC (BB/N000463/1). EEPJ was supported by BBSRC training grant BB/L502443/1 and Genus PLC. J.C. and C.I.R were supported by INSERM (*Institut National pour la Santé et la Recherche Médicale*). Their research was funded by ANR (*Agence Nationale pour la Recherche*; ANR-12–JSV2-0005– 01 and ANR-17-CE12-0004-01 to J.C).

## Authors’ contributions

Conceptualisation, PJIE; Methodology, BMS, PJIE; Software and Validation, BMS, CCR, PJIE; Investigation, CCR, BMS, EJ, DD, CP, GS, CIR; Data Curation and Formal Analysis, CCR, BMS, PJIE; Visualisation, BMS, PJIE; Supervision and Project Administration, PJIE, NA, JC; Writing - Original Draft, PJIE, BMS, CCR; Writing - Review and Editing, PJIE, BMS, CCR, GS; Resources, JC; Funding Acquisition, PJIE, NA, JC. All authors gave final approval for publication.

## Methods

### Mouse breeding

All animal procedures were in accordance with the United Kingdom Animal Scientific Procedures Act 1986 and were subject to local ethical review in UK and France (*Comite d’Ethique pour l’Experimentation Animale*, *Universite Paris Descartes*). Animals on MF1 strain background were bred on Home Office licenses 80/2451 and 70/8925, held by PE. These strains were originally sourced from colonies developed by Dr Paul Burgoyne (NIMR, Mill Hill, London), and were subsequently bred at Cambridge University Central Biomedical Services, or on contract by Charles River Laboratories (Manston, Kent, UK). Animals on C57Bl6/N background were bred at the Cochin Institute animal facility (licence held by JC, registration number CEEA34.JC.114.12). These strains were originally sourced from colonies developed by Dr Paul Burgoyne (NIMR, Mill Hill, London) and were subsequently backcrossed to C57Bl6/N animals at the Cochin Institute animal facility (to reach >95% of C57Bl6/N background). On both backgrounds, all males studied carry an RIII-derived Y chromosome. This provides the appropriate comparison for the ⅔ Yq deletion, which arose on a RIII background. Except when specified otherwise, breeding animals were housed as pairs or trios, and progeny housed singly or in small groups. Animals used in this study were sacrificed via CO_2_ followed by cervical dislocation (motility experiments) or cervical dislocation only (morphology experiments) and tissues collected post mortem for analysis.

### Superovulation and IVF

For the IVF work, and for *in vivo* experiments requiring superovulation, 5-6 week old females were induced to superovulate with injections of 5 IU PMSG (NVS Cat no. 859448) and 5 IU hCG (NVS Cat no.804745) in 100-150 µl of PBS, given 48 h apart. Donor males were sacrificed by cervical dislocation. One cauda epididymis was dissected and transferred to a 250 µl pre-equilibrated (37°C, 5% CO_2_) droplet of HTF media (Quinn et al., 1985) supplemented with 5% BSA, under embryo safe mineral oil (Sigma). The cauda was opened with a scalpel and the sperm allowed to swim out and capacitate for 1 hr at 37°C, 5% CO_2_. Next, superovulated females (4 for each cumulus-on and cumulus-off experimental sample) were sacrificed. Cumulus/oocyte complexes were retrieved from the ampulla of the oviduct and transferred either into a 500µl pre-equilibrated droplet of either HTF media or HTF media with hyaluronidase (300 µg/ml, Sigma H4272) and incubated for 3 minutes. The decumulated oocytes were rinsed briefly through a second HTF droplet and transferred to a final 500µl droplet of HTF media without hyaluronidase. 3µl of capacitated sperm were added to each fertilisation droplet and incubated for 4 hrs to allow fertilisation. After fertilisation, embryos were rinsed briefly through a fresh HTF droplet and transferred to culture wells containing 1ml KSOM media + 3mg/ml BSA for overnight incubation. The following morning, cleavage-stage embryos were transferred to KSOM without BSA and cultured to blastocyst stage for GFP scoring. Total fertilisation rates were 78.4% for cumulus-on oocytes and 82.5% for decumulated oocytes (not significant). Progression from 2-cell to blastocyst stages was 95.7% for cumulus-on versus 89.8% for cumulus-off groups (p = 0.013, binomial test for difference of proportions).

### Embryo recovery and timed mating experiments

For the 2-cell embryo recovery experiment, superovulated females were housed with males overnight (2 per male tested), and checked for plugs the following morning. Males were then re-mated to new superovulated females at intervals of 3/14/3 days following the first mating. This allowed collection of replicate data for long and short mating intervals from each male. Females were sacrificed at 1.5 days *post coitus* and 2-cell embryos cultured to the blastocyst stage to score GFP expression. 93-97% of recovered 2-cell embryos progressed to blastocyst stage. The difference in progression between genotypes was non-significant, and also too small in magnitude to explain the sex ratio skew.

For timed mating experiments looking at the effects of female background, estrus cycle progression was staged via visual inspection, females in estrus were housed with males overnight, and checked daily until plugged. Each male was mated successively at 1-week intervals to 4 different females: 2 from the XY^RIII^ colony and two from the XY^RIII^qdel colony. This allowed replicate data to be collected for both maternal backgrounds for each male tested. Once plugged, females were boxed out, sacrificed mid-gestation, and the number of green (XX) versus non-fluorescent (XY) embryos counted. The number of reabsorbing embryos was counted and found to be independent of both male and female genotypes.

### Sperm collection and fixation for morphology analysis

Mice were sacrificed by cervical dislocation and sperm collected from cauda epididymis and vas deferens as previously described [44]. The sperm samples were rinsed in 3 washes of 1xPBS and fixed in 3:1 methanol:acetic acid. **Table S1** shows details of the sperm samples analysed in the work reported here. Initial experiments used pooled samples, subsequent experiments analysed individual males. No differences were observed in summary statistics from pooled versus individual samples.

### Test tube swim-up motility fractionation

2 caudae epidydimides from freshly dissected males were transferred to 200µl pre-equilibrated (37°C, 5% CO_2_) HTF droplets under embryo safe mineral oil (Sigma), opened with a scalpel and the sperm allowed to swim out and capacitate for 1 hr at 37°C, 5% CO_2_. Following capacitation, general motility was checked under a light microscope before carefully transferring 50µl sperm suspension to the bottom of a test tube (Greiner Bio-One #120160) with 3ml of pre-equilibrated (37°C, 5% CO□) HTF media. Sperm were allowed to swim up through the overlying column of media for 30 minutes, and then successive 0.5ml aliquots were carefully pipetted from the top of the liquid meniscus, yielding six successive fractions from different swim depths.

In one experiment (Figure 5, n = 9 XY^RIII^, 10 XY^RIII^qdel males, MF1 background), these fractions were immediately spun down onto glass slides, processed for X/Y FISH, and counted blind. For X/Y counting, the cell-containing area of the slide was systematically scanned at high magnification (100x objective), and all observed cells scored for X/Y status. Counting was continued until at least 400 cells had been counted for each slide stained, or until all cells on the slide had been counted if fewer than 400 were present in any given fraction. The data were modelled using a beta regression using a logit link, with X proportion explained as an interaction of strain, live/dead state, and segment using the *betareg* R package, version 3.1.1 [72], and likelihood ratio testing was performed using the *rcompanion* package, version 2.1.1 [73].

In another experiment (**Figure S3**, n = 5 XY^RIII^ and 5 XY^RIII^qdel males), the top and bottom fractions only were stained using 200nM MitoTracker Red (M7512, Life Technologies) for 30 minutes at 37°C, 5% CO_2_, and then fixed as described above. Sperm were spun down and imaged at lower magnification (x60 objective) to allow visualisation of the whole midpiece. 5 males were analysed per genotype, and 55 images taken per fraction per male, each image containing 1-3 cells. Consistent image exposure times were used to allow accurate relative quantification of fluorescence intensity. Midpiece length and average staining intensity were measured using ImageJ.

### Image capturing and FISH for sperm morphological comparisons

In order to compare X and Y-bearing sperm we performed a capture / recapture analysis in which we first imaged sperm nuclei using DAPI fluorescence microscopy, then subjected them to FISH labelling and re-imaged them to determine their X/Y status. This was necessary because the chromatin swelling step required for FISH probe penetration distorts the detailed morphology of the sperm head (**Figure S1**).

Fixed sperm were dropped onto a poly-L-lysine coated slides, air dried, stained using Vectorshield Antifade Mounting medium with DAPI (Vector Laboratories, Peterborough, UK), and covered with 22×50mm cover slips. Slides were imaged on an Olympus BX61 epifluorescence microscope equipped with cooled CCDs and appropriate filters, and a motorised stage (Prior Scientific, Cambridge, UK). Images were captured using SmartCapture 3 and exported in TIFF format for downstream analyses. XYZ coordinates were recorded for each image taken to allow re-identification of cells. Slides were washed in 2xSSC for 2×10 minutes to remove mounting medium, and dehydrated through an ethanol series (70%, 80%, 100%, 2mins at RT). FISH was performed as previously described [45] using X and Y chromosome paints (Cytocell, Cambridge, UK), and images were captured as described above. The saved slide positions were used to perform automated batch capture of the previously imaged nuclei.

### Sperm morphological analysis

Morphology analysis was performed using the ImageJ plugin ‘Nuclear Morphology Analysis’ [44], version 1.15.1, available at https://bitbucket.org/bmskinner/nuclear_morphology/wiki/Home. In addition to the previouslY-described measurements, we developed a simple user interface to allow FISH images and pre-FISH images to be displayed side by side. This allowed simple manual sorting of pre-FISH sperm images into groups of X-bearing versus Y-bearing sperm to allow comparison. Data were exported for further processing in R 3.5.1 [74]. Summary statistics were calculated per sample and per strain for each chromosome, with errors propagated by quadrature. Two-dimensional Barnes-Hut tSNE [75] was run on angle profiles using *Rtsne*, version 0.15 [76] (perplexity=100, max_iter=1000) and a consistent seed. Parameters were chosen following confirmation of plot consistency across a range of values. Hierarchical clustering was performed by agglomerative nesting [77] using Ward’s method via the *cluster* package, version 2.0.7-1 [78]. Ward’s method was selected as giving the highest agglomerative coefficient. Dendrograms were cut, with cluster number determined following visual inspection, and cell cluster membership was imported back into Nuclear Morphology Analysis to allow consensus nucleus building. FISH images from males on the C57BL6 background were further used to assess nuclear organisation via dynamic signal warping, as previously described [45].

