## Supplemental Information for "Differential sperm motility mediates the sex ratio drive shaping mouse sex chromosome evolution"

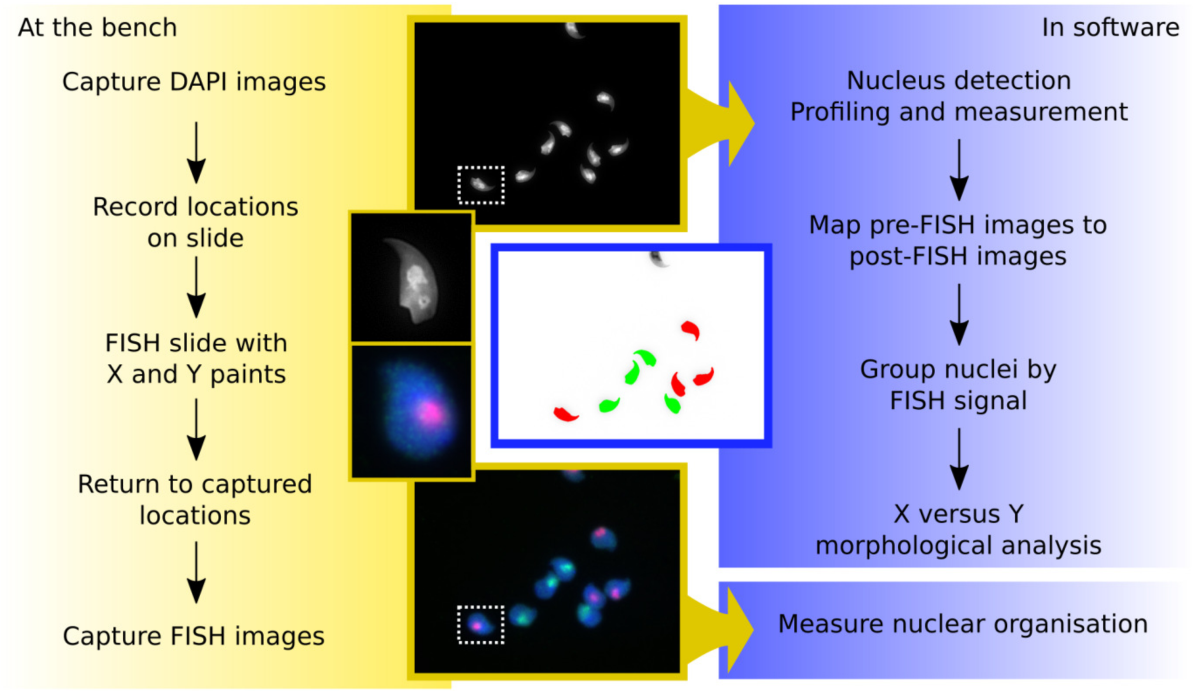


**Figure S1, related to FISH image capture methods.** Flowchart showing the capture-recapture protocol. The extensive swelling required for FISH probes to penetrate the nucleus (inset nuclei show before and after FISH) requires storing slide coordinates and imaging the slide before and after FISH for morphology analysis and subsequent XY grouping. A software user interface allows simple assignment of pre-FISH nuclei based on post-FISH images.

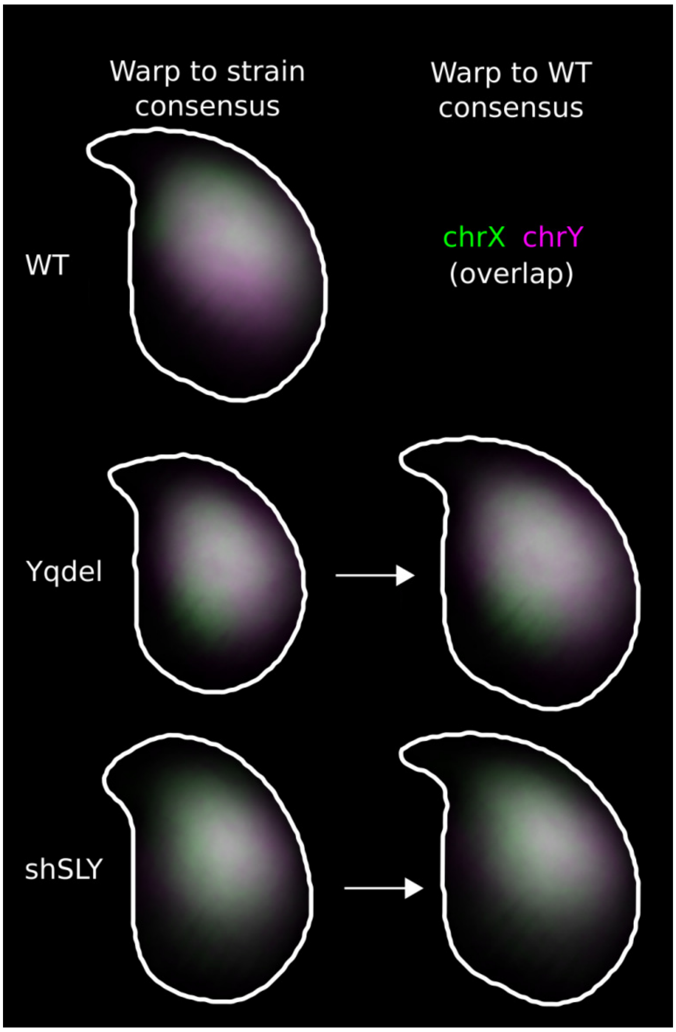


**Figure S2, related to Figure 4.** Chromosome territories for X and Y are sub-acrosomal in C57Bl6, and strongly overlap in all genotypes analysed (MS-SSIM* 0.86-0.94). Following warping to a WT template, the location of the X and Y chromosomes in sperm from Yqdel and shSLY males closely resembles their locations in wild type C57Bl6 sperm (MS-SSIM* 0.82-0.86). This indicates morphological changes are external to the nucleus, and do not involve other chromatin reorganisation.

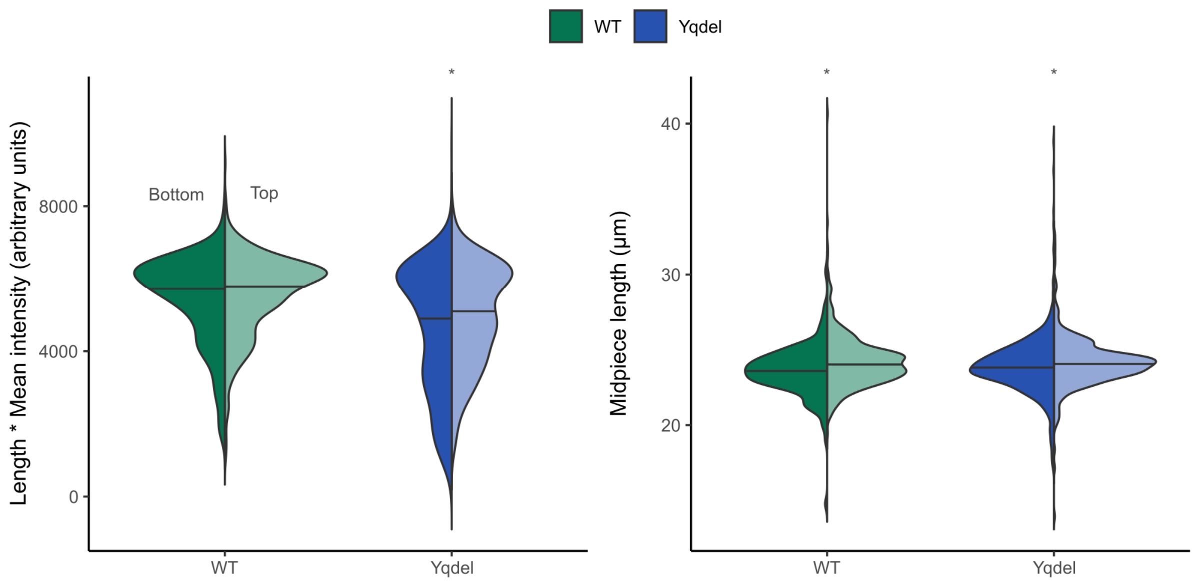


**Figure S3, related to Figure 5.** Midpiece length multiplied by mean signal is a proxy for total mitochondrial activity. Horizontal lines show sample medians. A two-sample two-tailed Kolmogorov-Smirnov test detects no difference between top and bottom for multiplied values in wild type $XY^{RIII}$ (p=0.73) and a significant difference in $XY^{RIII}$qdel (p=7.15E-05). Midpiece length is a proxy for mitochondrial count, and is slightly higher in top fractions in both $XY^{RIII}$ (p=0.0004) and $XY^{RIII}$qdel (p=0.006).

| Type | Sample | Number of animals | Ages (weeks) | n cells | X | Y | Average X area | Average Y area | Y/X area ratio | Area difference | S.E.M. | Volume difference | S.E.M. |
|---|---|---|---|---|---|---|---|---|---|---|---|---|---|
| MF1 WT | WT P1 | 8 | 13 - 16 | 1219 | 634 | 585 | 20.78 | 20.56 | 99.0% | 1.0% | 0.3% | 1.6% | 0.5% |
| | WT P2 | 8 | 10 - 14 | 777 | 426 | 351 | 20.37 | 20.05 | 98.4% | 1.6% | 0.3% | 2.3% | 0.5% |
| | WT I1 | 1 | 10.9 | 1524 | 777 | 747 | 20.93 | 20.73 | 99.0% | 1.0% | 0.2% | 1.4% | 0.3% |
| | WT I2 | 1 | 29.7 | 305 | 150 | 155 | 20.76 | 20.59 | 99.2% | 0.8% | 0.6% | 1.2% | 0.9% |
| | WT I3 | 1 | 26.1 | 413 | 210 | 203 | 20.28 | 19.88 | 98.1% | 1.9% | 0.5% | 2.9% | 0.8% |
| | *Average* | | | | | | *20.62* | *20.36* | *98.7%* | *1.3%* | *0.2%* | *1.9%* | *0.3%* |
| | **Aggregate** | | | 4238 | 2197 | 2041 | **20.70** | **20.47** | **98.9%** | **1.1%** | **0.2%** | **1.7%** | **0.2%** |
| MF1 Yqdel | Yqdel P1 | 5 | 11 - 14.5 | 904 | 386 | 518 | 18.50 | 18.02 | 97.4% | 2.6% | 0.4% | 3.8% | 0.5% |
| | Yqdel P2 | 6 | 9.3 | 308 | 119 | 189 | 18.93 | 18.03 | 95.3% | 4.7% | 0.6% | 7.0% | 0.9% |
| | Yqdel I1 | 1 | 11 | 860 | 440 | 420 | 19.76 | 19.35 | 97.9% | 2.1% | 0.3% | 3.1% | 0.5% |
| | Yqdel I2 | 1 | 11 | 262 | 136 | 126 | 18.79 | 18.35 | 97.7% | 2.3% | 0.6% | 3.5% | 1.0% |
| | Yqdel I3 | 1 | 11 | 574 | 279 | 295 | 18.93 | 18.58 | 98.1% | 1.9% | 0.3% | 2.8% | 0.5% |
| | Yqdel I4 | 1 | 31.3 | 177 | 91 | 86 | 19.31 | 18.55 | 96.1% | 3.9% | 0.9% | 5.8% | 1.3% |
| | Yqdel I5 | 1 | | 187 | 94 | 93 | 18.44 | 17.94 | 97.3% | 2.7% | 0.7% | 4.0% | 1.0% |

| | | | | | | | | | | | | | |
|---|---|---|---|---|---|---|---|---|---|---|---|---|---|
| | *Average* | | | | | | *18.95* | *18.40* | *97.1%* | *2.9%* | *0.4%* | *4.3%* | *0.5%* |
| | **Aggregate** | | | 3419 | 1638 | 1781 | **19.01** | **18.49** | **97.3%** | **2.7%** | **0.2%** | **4.1%** | **0.3%** |
| **C57Bl6 WT** | WT 1 | 1 | 24.6 | 1541 | 784 | 757 | 20.30 | 20.01 | 98.6% | 1.4% | 0.3% | 2.2% | 0.5% |
| | WT 2 | 1 | 24.6 | 1235 | 608 | 627 | 20.11 | 19.95 | 99.2% | 0.8% | 0.3% | 1.2% | 0.5% |
| | *Average* | | | | | | *20.21* | *19.98* | *98.9%* | *1.1%* | *0.8%* | *1.7%* | *1.1%* |
| | **Aggregate** | | | 2776 | 1392 | 1384 | **20.22** | **19.98** | **98.8%** | **1.2%** | **0.2%** | **1.8%** | **0.3%** |
| **C57Bl6 Yqdel** | Yqdel 1 | 1 | 14.6 | 1102 | 566 | 536 | 17.94 | 17.47 | 97.4% | 2.6% | 0.4% | 3.9% | 0.5% |
| | Yqdel 2 | 1 | 14.6 | 1101 | 594 | 507 | 17.87 | 17.36 | 97.2% | 2.8% | 0.3% | 4.2% | 0.5% |
| | Yqdel 3 | 1 | 11.4 | 690 | 344 | 346 | 17.60 | 17.14 | 97.4% | 2.6% | 0.4% | 3.9% | 0.6% |
| | Yqdel 4 | 1 | 11.4 | 735 | 355 | 380 | 17.04 | 16.62 | 97.5% | 2.5% | 0.4% | 3.7% | 0.6% |
| | *Average* | | | | | | *17.61* | *17.15* | *97.4%* | *2.6%* | *0.1%* | *3.9%* | *0.1%* |
| | **Aggregate** | | | 3628 | 1859 | 1769 | **17.68** | **17.19** | **97.2%** | **2.8%** | **0.2%** | **4.1%** | **0.3%** |
| **C57Bl6 shSLY** | shSLY 1 | 1 | 14.9 | 1149 | 587 | 562 | 17.89 | 17.65 | 98.6% | 1.4% | 0.6% | 2.1% | 0.9% |
| | shSLY 2 | 1 | 24.6 | 1261 | 653 | 608 | 17.84 | 17.65 | 98.9% | 1.1% | 0.5% | 1.6% | 0.7% |
| | shSLY 3 | 1 | 24.6 | 1368 | 666 | 702 | 17.99 | 17.81 | 99.0% | 1.0% | 0.5% | 1.4% | 0.7% |

| | | | | | | | | | | | | | |
|---|---|---|---|---|---|---|---|---|---|---|---|---|---|
| | shSLY 4 | 1 | 24.6 | 1160 | 575 | 585 | 18.80 | 18.53 | 98.6% | 1.4% | 0.6% | 2.2% | 0.9% |
| | *Average* | | | | | | *18.13* | *17.91* | *98.8%* | *1.2%* | *0.1%* | *1.8%* | *0.2%* |
| | **Aggregate** | | | 4938 | 2481 | 2457 | **18.11** | **17.90** | **98.8%** | **1.2%** | **0.3%** | **1.7%** | **0.4%** |

**Table S1, related to Figure 2 and Figure 3.** Details of the individual samples analysed, and the measured size differences in X- and Y-bearing sperm. S.E.M, standard error of the mean. The volume difference was calculated from area, assuming an equivalent reduction in sperm thickness.

| | Estimate | Std. Error | z value | Pr(>\|z\|) |
|---|---|---|---|---|
| (Intercept) | 0.015374 | 0.018551 | 0.829 | 0.407 |
| Segment | -0.010992 | 0.004764 | -2.307 | 0.021 |
| Strain alone | 0.038406 | 0.026236 | 1.464 | 0.143 |
| Live/dead state | -0.142011 | 0.024665 | -5.758 | 8.53E-09 |
| Segment:Strain | -0.001866 | 0.006737 | -0.277 | 0.782 |
| Segment:State | 0.032999 | 0.006333 | 5.211 | 1.88E-07 |
| Strain:State | 0.231879 | 0.034883 | 6.647 | 2.98E-11 |
| Segment:Strain:State | -0.049502 | 0.008956 | -5.527 | 3.25E-08 |

**Table S2, related to Figure 5**. Summary of the beta regression on swim-up data, showing the significance of each parameter and their interactions. The pseudo $R^2$ was 0.87.
